# An *RHD6*-directed gene network sustains root hair growth under combined low phosphate and salt stress in Arabidopsis

**DOI:** 10.64898/2026.11.25.769266

**Authors:** Miguel Angel Ibeas, Jorge Pérez-Díaz, Gabriel Padilla, Patricio Tapia-Reyes, Fiorella Garcia, Hernan Salinas-Grenet, Romina Acha, Claudio Meneses, José Manuel Estevez

## Abstract

Phosphate deficiency and salinity can coincide, but how roots integrate them is poorly understood. Root hairs (RHs) elongate under low inorganic phosphate and are inhibited by salt. Screening β-estradiol-inducible TRANSPLANTA lines for 18 RH-expressed transcription factors (TFs) in Arabidopsis (*Arabidopsis thaliana*), we found that induction of *ROOT HAIR DEFECTIVE 6* (*RHD6*) maintained RH growth under combined low phosphate (5 µM) and salt (100 mM NaCl), which inhibited wild-type RH growth. RNA sequencing of *RHD6*-induced roots under dual stress identified 1,811 differentially expressed genes. Up-regulated genes were enriched for RH differentiation, tip growth and cell wall organization; down-regulated genes for phenylpropanoid and suberin metabolism, hydrogen peroxide catabolism, photosynthesis and responses to wounding and jasmonic acid. Promoter motif scanning combined with ensemble co-expression analysis defined a core network of 396 candidate direct RHD6 targets, including 31 TFs. The TFs with the largest expression changes were *HEC3*, *HEC1* and *EIL4*. In T-DNA insertion mutants, dual stress did not shorten *hec3* RHs; *eil4* and *hec1* had shorter RHs on control medium, but their relative inhibition did not differ significantly from the wild type. These data provide a resource of RHD6-dependent genes and identify HEC3 as a modulator of the RH response to combined stress.

**Highlight:** Inducing the transcription factor RHD6 keeps Arabidopsis root hairs growing under combined phosphate deficiency and salinity; loss of its candidate target HEC3 makes root hairs insensitive to this stress.

## Introduction

Root hairs (RHs) are tubular outgrowths of root epidermal cells that enlarge the surface available for the uptake of water and nutrients (Gilroy and Jones, 2000). RHs improve phosphorus acquisition, including in soils that restrict root growth (Haling *et al*., 2013; Miguel *et al*., 2015), and their growth is highly responsive to the environment. Low availability of inorganic phosphate (Pi) stimulates RH elongation (Bates and Lynch, 1996; Datta *et al*., 2015; Bhosale *et al*., 2018), whereas salt reduces the number and length of RHs (Wang *et al*., 2008; Jin *et al*., 2023). In a meta-analysis, only 82 of 1,644 RH-specific genes responded to both Pi starvation and salt (Ibeas *et al*., 2024). Pi starvation and salinity also interact in shaping root system architecture (Kawa *et al*., 2016), but how RHs respond when both stresses are imposed together is not known.

In Arabidopsis (*Arabidopsis thaliana*), RH development depends on the class I RSL basic helix– loop–helix (bHLH) transcription factors (TFs) ROOT HAIR DEFECTIVE 6 (RHD6) and RHD6-LIKE 1 (RSL1) (Masucci and Schiefelbein, 1994; Menand *et al*., 2007). Downstream, the class II factor RSL4 determines final RH size (Yi *et al*., 2010; Datta *et al*., 2015) and binds the RH-specific *cis*-element of genes needed for RH morphogenesis (Hwang *et al*., 2017). The function of class I *RSL* genes is conserved among land plants (Menand *et al*., 2007; Kim and Dolan, 2016; Proust *et al*., 2016), and *RSL* genes are associated with RH length in wheat and rice (Han *et al*., 2016, 2017; Moon *et al*., 2019). Hormonal and environmental signals converge on this cascade. Auxin produced in response to low Pi or low nitrogen induces *RSL2* and *RSL4* or the RHD6–LRL3 module through AUXIN RESPONSE FACTORs (Mangano *et al*., 2017; Bhosale *et al*., 2018; Jia *et al*., 2023), PHR1 activates *RSL2* under low Pi (Yang *et al*., 2026), and ETHYLENE INSENSITIVE 3 (EIN3) interacts with RHD6 to co-activate *RSL4* (Feng *et al*., 2017). Circadian-clock PRR proteins, of which PRR5 binds RHD6, promote RH development, also under low Pi or salt (Bai *et al*., 2026). Conversely, ZINC FINGER PROTEIN 1, acting downstream of GLABRA2, represses *RHD6*, *RSL2* and *RSL4* (G. Han *et al*., 2020), JAZ proteins bind RHD6 and inhibit its activity (X. Han *et al*., 2020), MYB30 represses *RSL4* (Xiao *et al*., 2021) and, under salt stress, ABRE-binding factors (ABFs) bind RHD6 and suppress its transcriptional activity (Jin *et al*., 2023). *RHD6* transcription is in turn enhanced at low temperature by the long non-coding RNA *APOLO* and WRKY42 (Moison *et al*., 2021). Genes that depend on RHD6 have been catalogued in plants that lack it (Bruex *et al*., 2012; Feng *et al*., 2017; Zeng *et al*., 2023), but the genes that respond to an increase in RHD6 dosage have not been defined, particularly under combined stresses.

Here, we used the β-estradiol-inducible TRANSPLANTA (TPT) collection (Coego *et al*., 2014) to ask which RH-expressed TFs can maintain RH growth under combined low Pi and salt stress. Induction of *RHD6* was sufficient to do so. We combined RNA sequencing (RNA-seq) of *RHD6*-induced roots with promoter motif scanning and ensemble co-expression analysis to delineate a core set of candidate direct RHD6 targets, used T-DNA insertion mutants to test the three TFs of this set with the largest expression changes, HEC1, HEC3 and EIL4, and examined them in public single-stress datasets.

## Materials and methods

### Plant material and growth conditions

*Arabidopsis thaliana* (L.) Heynh. accession Columbia-0 (Col-0) was used as the wild type. The β-estradiol-inducible TPT lines carry TF coding sequences in the pER8GW vector (Zuo *et al*., 2000; Papdi *et al*., 2008; Coego *et al*., 2014); the 18 lines screened are listed in Dataset S1. Two independent *RHD6* lines, TPT_1.66470.1G and TPT_1.66470.1I (hereafter TPT-RHD6 1G and 1I), were used in all further experiments. The T-DNA insertion lines *hec1* (GABI_297B10; *HEC1*, AT5G67060), *hec3* (SALK_005294; *HEC3*, AT5G09750) and *eil4* (SALKseq_034500.1; *EIL4*, AT5G10120) carry the insertion in the coding sequence. Homozygous plants were selected by PCR with gene-specific primers and the T-DNA left-border primers LBb1.3 (SALK lines) or o8474 (GABI-Kat line); insertion sites and primers are given in Dataset S1. Seeds were surface sterilized, stratified at 4 °C in darkness for 3 d and germinated on half-strength Murashige and Skoog medium (0.5× MS; Murashige and Skoog, 1962) with MES (Duchefa, The Netherlands) and 0.8% (w/v) plant agar (Duchefa) in 120 × 120 mm square plates (Deltalab, Spain). Plates were kept vertically in a growth chamber at 22 °C under continuous light (120 µmol m^−2^ s^−1^).

### β-Estradiol induction and stress treatments

After 7 d on 0.5× MS (5 d in one experiment, as indicated in the figure legends), seedlings were transferred for 3 d to fresh 0.5× MS (control) or to dual-stress medium containing 5 µM Pi and 100 mM NaCl, in each case with or without β-estradiol. β-Estradiol was used at 1 µM unless indicated otherwise; in the screen shown in Fig. 1A it was used at 15 µM, and the control medium contained 0.075% DMSO.

**Fig. 1.**
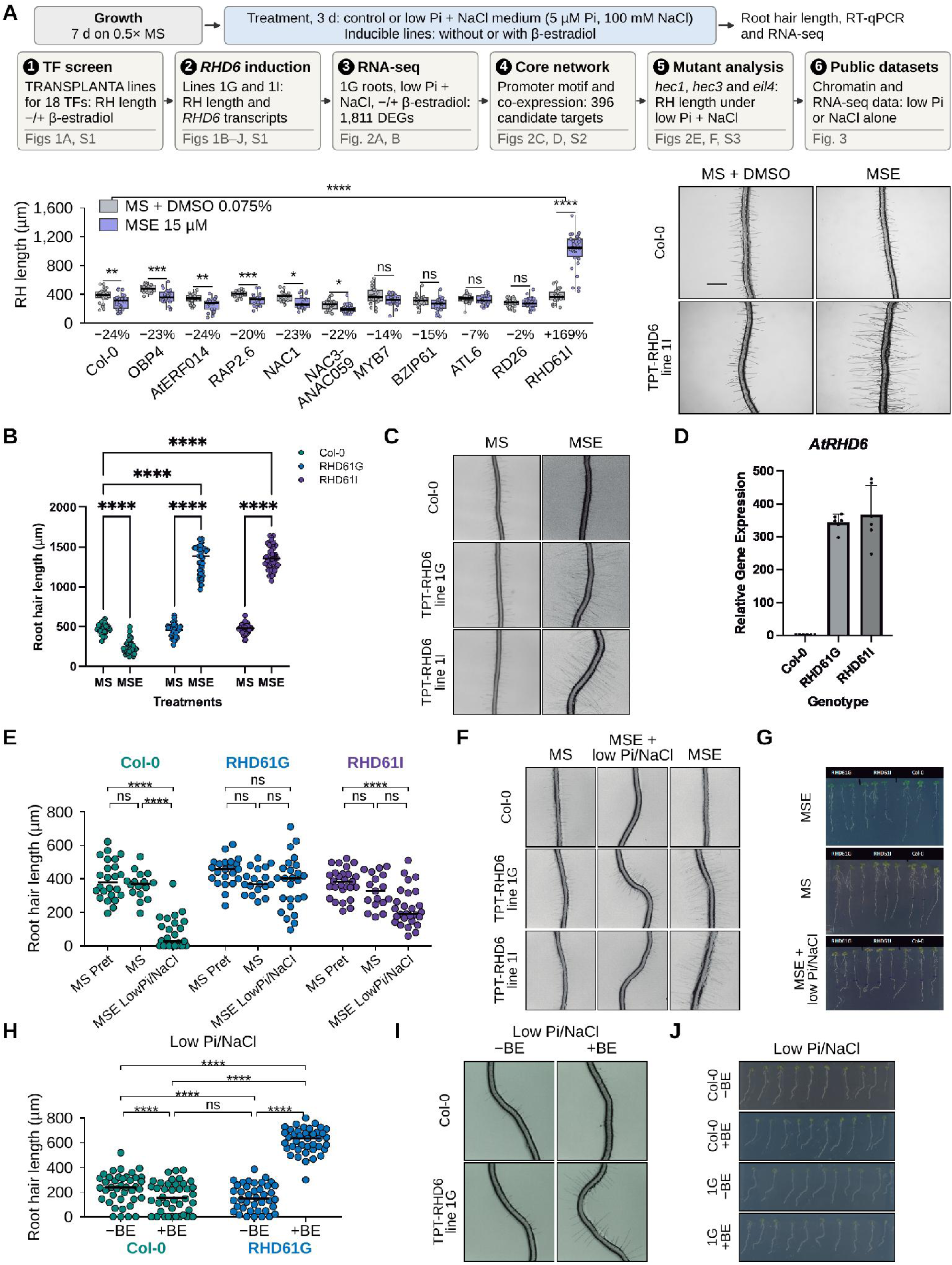
Induction of *RHD6* maintains root hair growth under combined low phosphate and salt stress. (A) Workflow of the study and root hair (RH) length in the TRANSPLANTA (TPT) screen. Top, workflow. Seedlings were grown for 7 d on 0.5× MS and transferred for 3 d to control medium or to medium with 5 µM Pi and 100 mM NaCl (low Pi + NaCl), with or without β-estradiol in the case of the inducible lines; the numbered boxes give the six steps of the study and the figures that report them. DEG, differentially expressed gene; TF, transcription factor. Bottom left, RH length in the screen. Col-0 and the TPT lines identified in Fig. S1B, including TPT-RHD6 1I (RHD61I), were grown for 7 d on 0.5× MS and transferred for 3 d to 0.5× MS with 0.075% DMSO (MS + DMSO) or with 15 µM β-estradiol (MSE). Each point is the mean length of the 10 longest RHs of one root; boxes show the median and interquartile range and whiskers the range. The numbers of roots (MS + DMSO/MSE) were 30/27, 24/25, 24/26, 27/26, 25/20, 23/29, 28/25, 22/28, 25/27, 29/30 and 28/30 for the lines in the order shown, from three experiments (two for NAC1 on MSE). Symbols above each line compare the two treatments (two-sided Welch’s *t*-test) and the bracket compares the response of TPT-RHD6 1I to β-estradiol with that of Col-0 (line × treatment interaction contrast); *P*-values were adjusted with the Holm method for all lines measured (Dataset S1): \*\*\*\**P*<0.0001; \*\*\**P*<0.001; \*\**P*<0.01; \**P*<0.05; ns, not significant. Numbers below the axis are the change in mean RH length with β-estradiol. Bottom right, representative roots of Col-0 and TPT-RHD6 1I; scale bar, 300 µm. (B) RH length of Col-0 and of two independent β-estradiol-inducible lines, TPT-RHD6 1G and 1I (RHD61G and RHD61I in the graphs), grown for 7 d on 0.5× MS and transferred for 3 d to 0.5× MS without (MS) or with β-estradiol (MSE). (C) Representative RHs of the plants in (B). (D) *RHD6* transcript levels relative to Col-0, measured by RT-qPCR in seedlings on MSE; bars are means ± SD and dots are individual values. (E) RH length before transfer (MS Pret; 7 d on 0.5× MS) and 3 d after transfer to 0.5× MS (MS) or to medium with 5 µM Pi and 100 mM NaCl plus β-estradiol (MSE LowPi/NaCl). (F) Representative RHs 3 d after transfer to the media indicated. (G) Seedlings 3 d after transfer. (H) RH length of Col-0 and TPT-RHD6 1G grown for 5 d on 0.5× MS and transferred for 3 d to medium with 5 µM Pi and 100 mM NaCl without (−BE) or with (+BE) 1 µM β-estradiol. (I) Representative RHs and (J) seedlings of the plants in (H). In (B), (E) and (H), each point is the mean length of the 10 longest RHs of one root and horizontal lines mark the group centre (the median in E and H); \*\*\*\**P*<0.0001; ns, not significant. The numbers of roots were 24/16/26 (Col-0), 22/18/25 (RHD61G) and 27/17/27 (RHD61I) for MS Pret/MS/MSE LowPi/NaCl in (E) and 40 per group in (H). Alt text: A flow diagram of the six steps of the study and box plots of root hair length in a screen of inducible transcription factor lines, in which only the RHD6 line forms longer root hairs after induction, followed by dot plots, micrographs of roots and photographs of seedlings showing that β-estradiol-induced *RHD6* lines form long root hairs on control medium and under low phosphate plus salt, whereas wild-type root hairs are strongly inhibited by the stress.

### Root hair phenotyping

Roots were imaged with a Leica EZ4 HD stereo microscope (Leica, Germany) and LAS EZ software. Only RHs that developed after transfer were analysed: the 10 longest RHs in the maturation zone of each root were measured with ImageJ (Schneider *et al*., 2012), and their mean was taken as the value for that root. Roots were pooled from three independent experiments.

### RT-qPCR

Total RNA was extracted from roots with the E.Z.N.A. Total RNA Kit (Omega Bio-tek, USA). One microgram of RNA was reverse transcribed with an oligo(dT)_20_ primer (Macrogen, South Korea) and M-MLV reverse transcriptase (Invitrogen, USA). The cDNA was diluted 3-fold, and 1 µl was amplified in 20 µl reactions containing 10 µl of Brilliant III Ultra-Fast SYBR Green Master Mix (Stratagene, USA) and 0.25 µM of each primer in an AriaMx Real-Time PCR System (Agilent, USA). Transcript levels were normalized to *TAFII15* (AT4G31720), whose transcript abundance changed only 1.2-fold (adjusted *P*=0.02) after *RHD6* induction in the RNA-seq dataset (Dataset S1), and are shown as means ± SD.

### RNA-seq and differential expression

TPT-RHD6 1G seedlings were grown for 7 d on 0.5× MS and transferred for 3 d to dual-stress medium with or without 1 µM β-estradiol; roots of three biological replicates per treatment were harvested. Libraries were sequenced as 151-bp paired-end reads, yielding 188–231 million reads per library (Dataset S1), and the data were deposited at the NCBI under BioProject accession number PRJNA1565760. Reads were trimmed with Trim Galore v0.6.10 (Krueger *et al*., 2023) using the options ‘--length 50 --paired -q 20’ and aligned to the TAIR10 genome with the Araport11 annotation (Cheng *et al*., 2017) using STAR v2.7.11b (Dobin *et al*., 2013) with default parameters; 72–82% of the reads mapped uniquely. Reads per gene were counted with featureCounts (Liao *et al*., 2014) from Subread v2.0.2 (Liao *et al*., 2019). Differential expression between induced and non-induced samples was tested with DESeq2 v1.49.1 (Love *et al*., 2014) for the 21,513 genes that passed its filters. Genes with an adjusted *P*-value (*P*_adj_) ≤0.01 and an absolute log_2_ fold change ≥1.5 were considered differentially expressed genes (DEGs). For the heatmap, DESeq2-normalized counts of the DEGs were converted to row *z*-scores and clustered by Euclidean distance with average linkage.

### Gene Ontology enrichment

Enrichment of Gene Ontology (GO) biological process terms was tested separately for up- and down-regulated DEGs with GOATOOLS v1.6.9 (Klopfenstein *et al*., 2018), using the GO release of 26 July 2026, the TAIR annotation file of 21 May 2026 and, as background, the 12,853 annotated genes among those tested by DESeq2. Annotations were propagated through *is_a* and *part_of* relations, and *P*-values from Fisher’s exact test were adjusted with the Benjamini–Hochberg method; terms with a false discovery rate (FDR) <0.01 were considered enriched.

### Promoter motifs, co-expression network and functional classes

The 3,000 bp upstream of each gene were scanned for the G-box (CACGTG) and for the 8-bp core (GGKCCKRW) of the GCG-box (GG[GT]CC[GT][GA][TA]C; Liu *et al*., 2015) with the ‘Statistical Motif Analysis in Promoter or Upstream Gene Sequences’ tool of TAIR (https://v2.arabidopsis.org/tools/bulk/motiffinder/index.jsp). The proportion of motif-containing promoters among DEGs and among the remaining tested genes was compared with Fisher’s exact test. Standardized co-expression strengths between DEGs were obtained from the Arabidopsis TEA-GCN ensemble co-expression network (Lim *et al*., 2026), kindly provided by its authors, and gene pairs with *z* ≥1.5 were considered highly co-expressed. The core gene regulatory network (GRN) comprised the DEGs that carry at least one motif and are highly co-expressed with *RHD6*. TFs were identified with PlantTFDB (Jin *et al*., 2017), and network genes were assigned to functional classes with ordered rules based on PlantTFDB, GO annotations and gene descriptions (Dataset S1). To evaluate the set of genes co-expressed with *HEC1*, *HEC3* and *EIL4*, its size was compared with that obtained for all triplets of network genes whose number of co-expression partners was within ±10 of that of each TF (2,964 triplets).

### Analysis of public datasets

All datasets were obtained from the NCBI Gene Expression Omnibus (Dataset S1). The 3-kb promoters of *HEC1*, *HEC3* and *EIL4* (TAIR10) were scanned on both strands for the G-box, the GCG-box core and the RH-specific *cis*-element bound by RSL4 (Hwang *et al*., 2017), and motif frequencies were obtained for the promoters of the 27,420 nuclear protein-coding genes. Chromatin accessibility (ATAC-seq), H3K27me3 (CUT&Tag) and nuclear RNA-seq data of root hair nuclei from Pi-replete plants and from plants deprived of Pi for 96 h were taken from GSE271993 (Holder and Deal, 2024, Preprint); because these data are mapped to the Col-PEK assembly, each locus was located by exact sequence match. Open-chromatin peaks (DNase-seq and FAIRE-seq) of control and NaCl-treated seedlings were from GSE145742 (Raxwal *et al*., 2020). Single-cell RNA-seq data of root tips under control, NaCl and abscisic acid treatments (GSE268881; Wang *et al*., 2026) were filtered and normalized with Scanpy v1.13 (Wolf *et al*., 2018), the six samples were integrated with Harmony (Korsunsky *et al*., 2019) and cells were clustered with the Leiden algorithm (Traag *et al*., 2019); clusters were assigned to cell types with marker genes, and expression was summarized as pseudobulk counts per million (CPM) and as the percentage of cells with a detected transcript. Bulk RNA-seq data of roots, root tips, sorted root cell types or root hair nuclei under low Pi (GSE131056, Silva-Navas *et al*., 2019; GSE128250, Blein *et al*., 2020; GSE148457, Ojeda-Rivera *et al*., 2020; GSE271932, Holder and Deal, 2024, Preprint; GSE287184, González-García *et al*., 2026) or NaCl (GSE195912, Augstein and Carlsbecker, 2022; GSE307070 and GSE270516, no linked publication) were converted to log_2_(CPM+1). The response to each stress was calculated for wild-type samples and compared with the response to *RHD6* induction by Spearman correlation across the DEGs and by Fisher’s exact test. For co-expression, genes with a mean of at least 1 CPM were standardized within each study, and Pearson correlations were combined across studies after Fisher z-transformation. These analyses are descriptive: the public experiments differ from ours in growth conditions, plant age and tissue, and none combines low Pi with NaCl. Filters, marker genes and all values are given in Dataset S1.

### Statistical analysis

In all RH experiments the unit of analysis was the root (one value per seedling; *n* is the number of roots) and the groups compared were independent. RH length data from the induction experiments in Fig. 1B, E, H and Fig. S1C were analysed with GraphPad Prism 8.0.1. For the T-DNA insertion mutants, values further than 1.5 times the interquartile range from the quartiles of their group were treated as outliers and excluded (one of 240 roots; Dataset S1), and control and dual-stress groups of each genotype were compared with two-sided Welch’s *t*-tests. Differences among genotypes in the response to stress were tested with a two-way ANOVA (genotype × condition) and with interaction contrasts against Col-0 that allow unequal variances (Welch–Satterthwaite approximation); 95% confidence intervals of the relative change in mean RH length were obtained by bootstrapping (20,000 resamples). Because the mutants differed from Col-0 in RH length on control medium, the relative change of each mutant was also compared with that of Col-0 as the ratio of the two stress-to-control ratios of mean RH length, tested on the log scale with delta-method standard errors and Welch–Satterthwaite degrees of freedom. The screen (Fig. 1A) was analysed in the same way: the two treatments were compared within each line with two-sided Welch’s *t*-tests and the response of each line to β-estradiol was compared with that of Col-0 with interaction contrasts, and *P*-values were adjusted with the Holm method (Dataset S1). These analyses were run in Python 3.13 with SciPy 1.18 and statsmodels 0.15.

## Results and discussion

### Induction of *RHD6* maintains root hair growth under combined low Pi and salt stress

The TPT collection was built to express each of 949 TFs from a β-estradiol-inducible promoter (Coego *et al*., 2014). We selected lines for 18 TFs that are expressed in RH cells according to cell type-resolved root transcriptomes (Brady *et al*., 2007; Denyer *et al*., 2019; Jean-Baptiste *et al*., 2019; Ryu *et al*., 2019; Shulse *et al*., 2019; Zhang *et al*., 2019) (Fig. S1A; Dataset S1). Seedlings grown for 7 d on 0.5× MS were transferred for 3 d to control medium or to medium with 5 µM Pi and 100 mM NaCl, with or without β-estradiol. On control medium, β-estradiol alone shortened RHs by 24% in Col-0 and by 20–24% in five of the TF lines shown, and had no significant effect in four others, whereas it increased RH length 2.7-fold in the *RHD6* line (from 375 to 1,007 µm), the only line whose response differed from that of Col-0 (Fig. 1A; Fig. S1B; Dataset S1). Among the lines tested, RH growth under dual stress was greatest in those expressing *RHD6*.

On 0.5× MS, β-estradiol increased RH length about 3-fold in two independent lines, TPT-RHD6 1G and 1I (from about 470 to about 1,400 µm; *P*<0.0001), in agreement with the RH proliferation reported for induced *RHD6* TPT lines (Coego *et al*., 2014), whereas it reduced RH length in Col-0 by about half (Fig. 1B, C). *RHD6* transcript levels in induced TPT lines were more than 300 times those of Col-0 (Fig. 1D). RH length of TPT-RHD6 1G was similar with 1, 5 and 7.5 µM β-estradiol, and 1 µM was used thereafter (Fig. S1C–E). Under dual stress with β-estradiol, RHs of Col-0 were almost completely inhibited (about 30 µm, compared with about 370 µm on 0.5× MS; *P*<0.0001), whereas those of TPT-RHD6 1G were as long as on control medium and those of line 1I were intermediate (about 190 µm) (Fig. 1E–G). In an independent experiment, RHs of induced TPT-RHD6 1G roots under dual stress were about four times longer than those of non-induced roots (about 640 versus 150 µm; *P*<0.0001), whereas in Col-0 β-estradiol reduced RH length from about 240 to about 150 µm (*P*<0.0001; Fig. 1H–J). As β-estradiol by itself shortens Col-0 RHs (Fig. 1A, B), the inhibition of Col-0 in these experiments combines the effects of the stress and of the inducer; the effect of the stress alone is quantified below. A high level of *RHD6* expression is therefore sufficient to maintain RH growth when Pi limitation and salinity are combined. Higher *RHD6* expression also restores RH growth under long-term mild heat (Du *et al*., 2025). Opposing inputs converge on the RSL cascade: low Pi induces *RSL2* and *RSL4* through auxin (Bhosale *et al*., 2018), whereas abscisic acid represses *RSL2* through OBP4 (Rymen *et al*., 2017). Salt was reported to reduce the long-RH phenotype of plants overexpressing *RHD6*, because ABFs bind RHD6 and suppress its transcriptional activity (Jin *et al*., 2023). As ABFs, like JAZ proteins, inhibit RHD6 by physical interaction, induced RHD6 levels may exceed what these repressors can neutralize.

### *RHD6* induction reinforces the root hair programme and lowers the expression of barrier, metabolic and stress-response genes

RNA-seq of TPT-RHD6 1G roots kept for 3 d under dual stress with or without β-estradiol identified 1,811 DEGs among 21,513 expressed genes, 750 up-regulated and 1,061 down-regulated (Fig. 2A; Dataset S1). *RHD6* itself increased 140-fold. No β-estradiol-treated Col-0 control was sequenced, so RHD6-independent effects of β-estradiol, which by itself shortened RHs in Col-0 (Fig. 1A, B, H), cannot be excluded. Up-regulated genes included the known downstream regulators *RSL2*, *RSL4* and *LRL3* and were enriched for trichoblast differentiation, RH elongation, tip growth and plant-type cell wall organization (Fig. 2B); they comprised several *EXTENSIN* genes, whose products are required for RH growth (Velasquez *et al*., 2011). Down-regulated genes were enriched for phenylpropanoid, suberin and terpenoid metabolism, assembly of the Casparian strip (*CASP1*– *CASP5*), hydrogen peroxide catabolism (18 class III peroxidase genes), photosynthesis, iron homeostasis (including *IRT1* and *FRO2*) and responses to wounding, jasmonic acid and water deprivation (Fig. 2B; Dataset S1). The term ‘response to salt stress’ was not enriched in either set. The up-regulated set fits the hierarchy defined genetically, in which class I RSL factors control the class II *RSL* genes (Yi *et al*., 2010; Pires *et al*., 2013); it includes genes that depend on RHD6 in the root epidermis, such as *LRX1* and *PRP3* (Bruex *et al*., 2012), and RSL4-activated effectors (Mangano *et al*., 2017) such as *RHD2* and the RH-specific *PRX01* and *PRX73* (Marzol *et al*., 2022). The down-regulated set, less expected for an activator of an epidermal programme, comprises suberin biosynthesis genes with four activators of suberin deposition, *MYB9*, *MYB107*, *MYB53* and *MYB93* (Lashbrooke *et al*., 2016; Shukla *et al*., 2021), and much of the Fe-acquisition machinery, with *IRT1* and *FRO2* reduced more than 200-fold together with their regulator *FIT* (Colangelo and Guerinot, 2004). Suberization responds to nutrient stress (Barberon *et al*., 2016), and suppression of photosynthesis genes in roots sustains root growth under Pi deficiency (Kang *et al*., 2014). These changes may reflect ectopic RHD6 activity, as the XVE transactivator is driven by a constitutive promoter (Zuo *et al*., 2000); sequestration of partners shared with other pathways, such as ABFs (Jin *et al*., 2023) or EIN3, which also binds FIT (Lingam *et al*., 2011; Feng *et al*., 2017); or an altered nutrient and water status caused by the longer RHs.

**Fig. 2.**
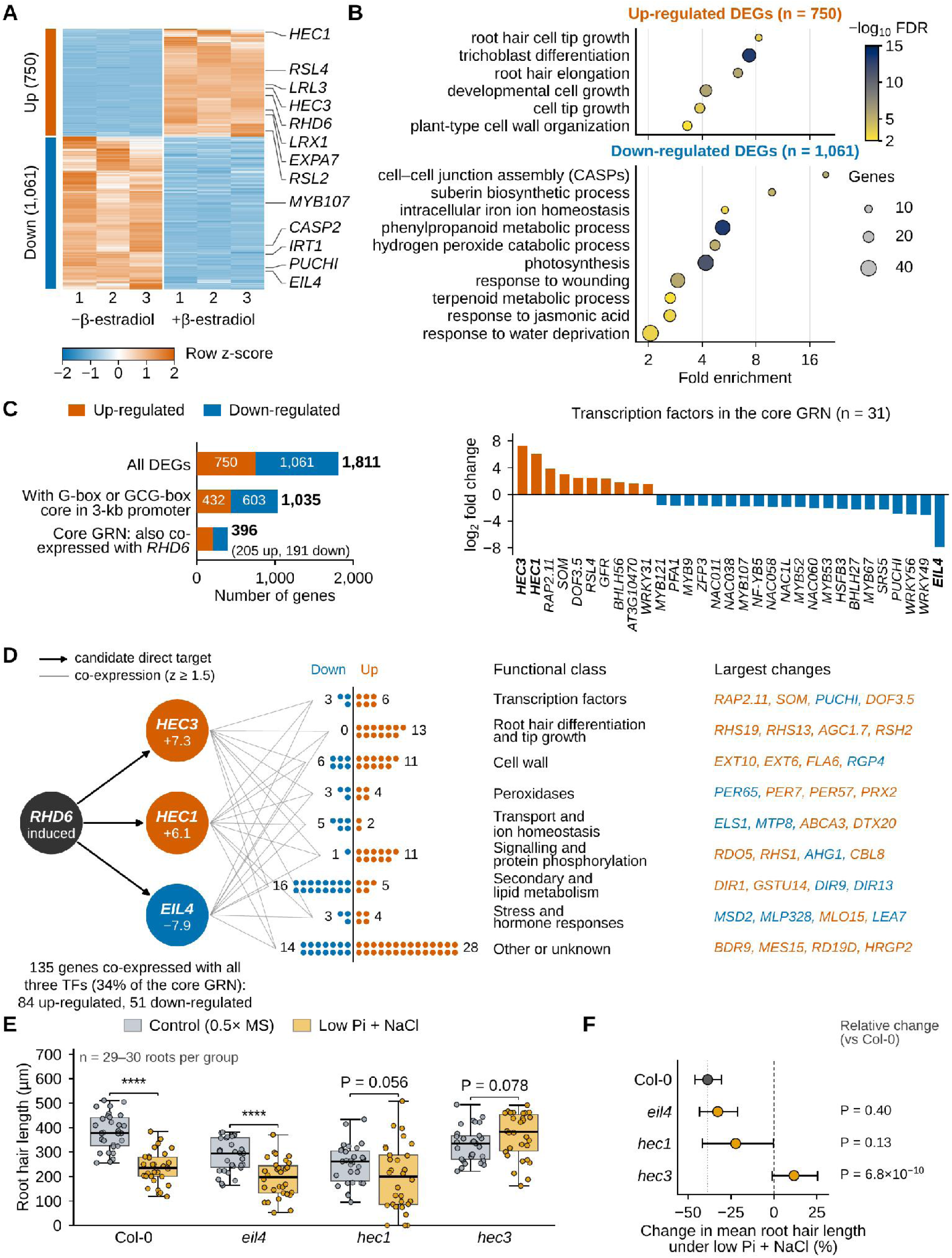
Transcriptome and core gene regulatory network of *RHD6*-induced roots under combined low phosphate and salt stress, and root hair response of *eil4*, *hec1* and *hec3* mutants. (A) Row *z*-scores of DESeq2-normalized counts for the 1,811 differentially expressed genes (DEGs) between TPT-RHD6 1G roots kept for 3 d on medium with 5 µM Pi and 100 mM NaCl without or with β-estradiol (three biological replicates each). Rows were clustered by Euclidean distance with average linkage; selected genes are indicated. (B) Representative Gene Ontology biological process terms enriched among up- and down-regulated DEGs (FDR <0.01; background, all genes tested); the complete results are given in Dataset S1. (C) Left, numbers of up- and down-regulated genes among all DEGs, among DEGs with a G-box or the GCG-box core in the 3-kb promoter and in the core network, which also requires co-expression with *RHD6* (*z* ≥1.5). Right, log_2_ fold change of the 31 transcription factor (TF) genes of the core network; the three with the largest changes are in bold. (D) The three prioritized TFs and the 135 core network genes co-expressed with all of them, grouped by functional class. Each dot is one gene (colours as in C; down-regulated genes to the left and up-regulated genes to the right of the line, with their numbers); arrows, candidate direct targets of RHD6; grey lines, co-expression (*z* ≥1.5); numbers, log_2_ fold change. The genes named are those with the largest changes in each class. (E) Root hair (RH) length of Col-0 and of homozygous T-DNA insertion mutants (Fig. S3) 3 d after transfer to control medium (0.5× MS) or to medium with 5 µM Pi and 100 mM NaCl. Each point is the mean of the 10 longest RHs of one root (*n*=30 roots per group; 29 for *hec3* on control medium after one outlier was removed, see Materials and methods); boxes show the median and interquartile range and whiskers the range. Control and stress were compared within each genotype with two-sided Welch’s *t*-tests: \*\*\*\**P*<0.0001; exact *P*-values are shown for the other comparisons. (F) Change in mean RH length under dual stress relative to the control (points) with bootstrap 95% confidence intervals (bars); the dotted line marks the change in Col-0. *P*-values compare the relative change of each mutant with that of Col-0 (ratio of the two stress-to-control ratios of mean RH length). The genotype × condition interaction in a two-way ANOVA was significant (*F*_3,231_=12.34, *P*<0.0001); the interaction contrasts on the absolute scale are given in Dataset S1. Alt text: Heatmap of 1,811 genes separating induced from non-induced samples, dot plot of enriched Gene Ontology terms, bar charts of gene numbers and of fold changes for 31 transcription factors, a diagram linking RHD6 to HEC3, HEC1 and EIL4 and to 135 shared co-expressed genes, and box plots showing that low phosphate plus salt shortens root hairs in the wild type and in *eil4*, less clearly in *hec1* and not in *hec3*, with a plot of the percentage change in each genotype.

### A core network of candidate direct RHD6 targets

To approximate the direct targets of RHD6, we searched 3-kb promoters for the G-box and the core of the GCG-box (Liu *et al*., 2015), because RHD6 binds *ALMT1* promoter fragments that carry these motifs (Cao *et al*., 2024). At least one motif was present in 1,035 DEGs (57%), the same proportion as among the other expressed genes (57%; *P*=0.86), so motif presence alone is not informative. We therefore retained the motif-containing DEGs that are also highly co-expressed with *RHD6* across public RNA-seq samples in the TEA-GCN ensemble network (Lim *et al*., 2026). This defined a core GRN of 396 candidate direct targets, 205 up-regulated and 191 down-regulated (Fig. 2C; Fig. S2; Dataset S1). It includes *RSL4*, the extensin genes *EXT6*, *EXT10* and *LRX1*, and the strongly repressed *IRT1*. Because the network rests on predicted motifs and co-expression, and the DNA-binding specificity of RHD6 has not been determined *in vitro*, these genes are candidates that await confirmation by chromatin immunoprecipitation. The approach is also conservative: *RSL2* and *LRL3*, which respond to RHD6, lack both motifs in the scanned region and are absent from the core GRN. Of the experimentally established targets, RHD6 and EIN3 directly co-activate *RSL4* (Feng *et al*., 2017), and in the response to aluminium RHD6 activates *ALMT1* and represses *RAE1* (Cao *et al*., 2024). *RSL4* and *ALMT1* belong to the core network, but *ALMT1* was repressed about 4-fold and *RAE1* did not respond. The effect of RHD6 on a target thus depends on the condition, as expected for a factor that acts with EIN3 and HY5 (Feng *et al*., 2017; Zhang *et al*., 2025). The network also contains 11 of the 19 *ROOT HAIR SPECIFIC* genes, whose promoters carry the RH-specific *cis*-element bound by RSL4 (Won *et al*., 2009; Hwang *et al*., 2017), so part of it probably responds to RHD6 through RSL4.

The core GRN contains 31 TF genes, 10 up-regulated and 21 down-regulated; the six *MYB* and five *NAC* genes among them are all down-regulated (Fig. 2C). The largest changes were those of the bHLH genes *HEC3* (log_2_ fold change 7.3) and *HEC1* (6.1) and of the *EIN3-LIKE* gene *EIL4* (−7.9), although *HEC1* and *EIL4* are expressed at low levels in whole roots (mean normalized counts ≤25 and ≤106; Dataset S1). In the co-expression network, these three TFs are linked to each other and share 135 co-expressed GRN genes (34% of the network; 84 up-regulated and 51 down-regulated), which include RH, cell wall and signalling genes (Fig. 2D). Degree-matched random triplets shared 91 ± 20 genes (mean ± SD; empirical *P*=0.016), but the difference was not significant when triplets were also required to be mutually co-expressed (*P*=0.11; Dataset S1). The network is dense, with 70% of all gene pairs connected, and the three TFs have fewer partners than the median gene (186–237 versus 299); they were therefore prioritized by the magnitude of their response to RHD6 and not by network centrality. As co-expression is undirected, it does not show that HEC1, HEC3 or EIL4 regulate the genes they share.

### Loss of *HEC3* abolishes the inhibition of root hair growth by dual stress

We next measured RH length in homozygous T-DNA insertion mutants of the three TFs (Fig. 2E, F; Fig. S3). In Col-0, dual stress reduced mean RH length by 39% (median 379 µm on control medium and 234 µm under stress; *P*<0.0001). The net effect of the combined stress on wild-type RHs is thus inhibitory, as if the response to salt prevailed over the stimulation of RH elongation by low Pi; lateral roots likewise prioritize the response to salt under this combination (Kawa *et al*., 2016). This inhibition is much weaker than that seen with β-estradiol (Fig. 1E), in line with the effect of the inducer; without it, Col-0 RHs under dual stress were about 230–240 µm long in both experiments (Figs 1H, 2E). In *hec3*, RH length did not decrease (+12%; *P*=0.08), unlike in Col-0 (relative change, *P*<0.0001): *hec3* RHs were shorter than those of Col-0 on control medium (median 336 µm; *P*=0.008) but much longer under stress (382 versus 234 µm). RHs of *eil4* and *hec1* were 25% and 34% shorter than those of Col-0 on control medium (both *P*<0.0001), and dual stress shortened them by 33% (*P*<0.0001) and 22% (*P*=0.06), respectively. Neither relative change differed significantly from that of Col-0 (*P*=0.40 and *P*=0.13), although the absolute decrease was smaller in both mutants (interaction *P*=0.04 and *P*=0.009). Thus HEC3 appears necessary for the inhibition of RH growth by combined low Pi and salt, whereas loss of EIL4 or HEC1 shortens RHs without detectably changing their relative response to stress.

The phenotype of *hec3* was unexpected. *HEC3* is among the genes most strongly induced by RHD6, yet its loss makes RH growth insensitive to the dual stress in plants that do not overexpress *RHD6*. HEC proteins are bHLH factors that act in gynoecium development, shoot meristem function and light signalling, modulate auxin and cytokinin responses (Gremski *et al*., 2007; Schuster *et al*., 2015; Gaillochet *et al*., 2018) and can inhibit other bHLH proteins by heterodimerization, as shown for HEC2 with PIF1 and PIF4 (Zhu *et al*., 2016; Lee *et al*., 2021). As PIFs activate *HEC* genes and HEC proteins then limit PIF activity, HEC3 induced by RHD6 may similarly restrain RSL-type bHLH factors, and thereby RH growth, under stress, which would explain the longer RHs of *hec3*; alternatively, HEC3 may act in the stress response independently of RHD6. Negative regulators are known in RH networks: GTL1 and DF1 repress *RSL4* (Shibata *et al*., 2022; Urzúa Lehuedé *et al*., 2025). EIL4 is a little-studied member of the EIN3 family. EIN3 accumulates under Pi starvation and salinity, interacts with RHD6 and promotes RH growth (Peng *et al*., 2014; Song *et al*., 2016; Feng *et al*., 2017); the shorter RHs of *eil4* and *hec1* suggest that EIL4 and HEC1 promote RH growth, so repression of *EIL4* would not explain the effect of *RHD6* induction. Testing these hypotheses will require combining each mutation with *RHD6* induction, higher-order *hec* mutants to address redundancy within the HEC clade, and independent alleles or complementation of each mutant.

### Public datasets place the network in the context of each single stress

We re-analysed public chromatin, single-cell and bulk RNA-seq data for low Pi or NaCl alone (Fig. 3; Dataset S1). The response to *RHD6* induction correlated positively with the response of whole roots to low Pi in two datasets (Spearman ρ=0.43 and 0.48 across DEGs; −0.11 to 0.20 in six others) and negatively with the response to NaCl in all six comparisons (ρ=−0.10 to −0.33; Fig. 3D). NaCl lowered *LRX1*, *PRP3* and *PRX73* transcripts in five of six comparisons, whereas *RHD6* transcripts changed at most 1.5-fold, also in trichoblasts (Fig. 3B, C). *RHD6* was co-expressed with *RSL2*, *LRL3* and these RH effector genes in low-Pi datasets (combined *r*=0.59–0.71) but not in NaCl datasets (*r*=−0.08 to 0.06; Fig. 3E). Salt thus uncouples the RH programme from *RHD6* transcript levels, as expected if ABFs inhibit RHD6 post-translationally (Jin *et al*., 2023), and *RHD6* induction restores it. *HEC1*, *HEC3* and *EIL4* were barely detectable in every public dataset and cell type (at most 4.1, 2.2 and 0.4 CPM, respectively) and carry high H3K27me3 in root hair nuclei (Fig. 3A–C); their co-expression with *RHD6* therefore probably arises in other tissues or conditions, and their response to induced RHD6 may be ectopic unless dual stress activates them in wild-type roots. The G-box of *HEC1* (−493 bp) and the GCG-box of *EIL4* (−1,188 bp), but not the *HEC3* motifs, lie in accessible chromatin of root hair nuclei with or without Pi (Fig. 3A).

**Fig. 3.**
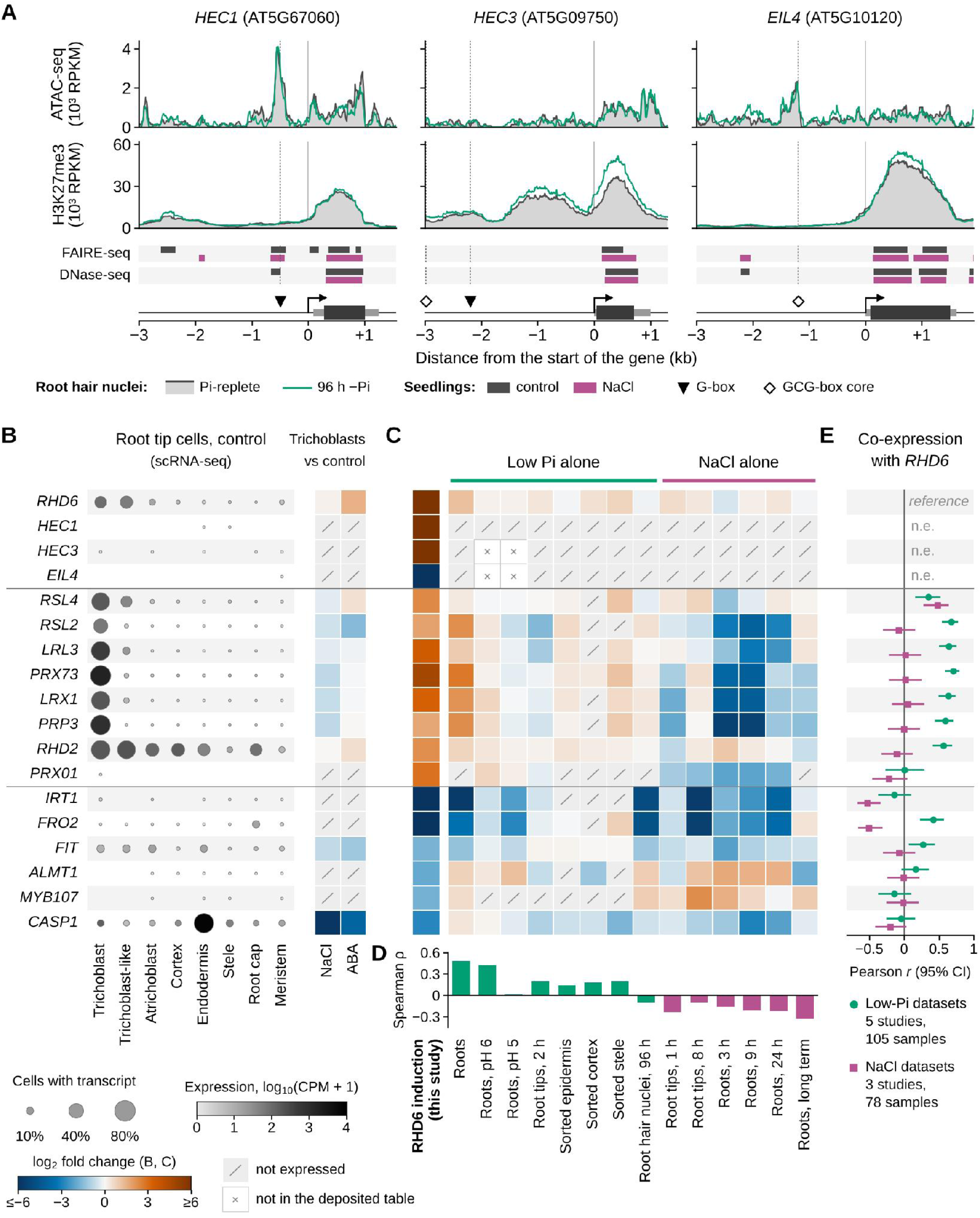
Public chromatin, single-cell and bulk transcriptome data for *RHD6*, *HEC1*, *HEC3*, *EIL4* and selected network genes under low phosphate or salt stress alone. (A) Promoters (3 kb) and transcribed regions of *HEC1*, *HEC3* and *EIL4*, drawn to the same scales. Top, ATAC-seq and H3K27me3 signal in root hair nuclei of Pi-replete plants and of plants deprived of Pi for 96 h (means of two and three replicates, respectively, in 10-bp bins; GSE271993; Holder and Deal, 2024, Preprint). Bottom, open-chromatin peaks of control and NaCl-treated seedlings (FAIRE-seq and DNase-seq; GSE145742; Raxwal *et al*., 2020), positions of the G-box and of the GCG-box core (dotted lines), and gene models (dark boxes, coding sequence; arrow, start of the gene). (B–E) Each row is one gene. (B) Expression in root tip cells (single-cell RNA-seq; GSE268881; Wang *et al*., 2026). Left, control cells grouped by cell class; dot size, percentage of cells with a detected transcript; shading, pseudobulk expression in counts per million (CPM); clusters without enrichment of a marker set (10% of the cells) are not shown. Right, log_2_ fold change in trichoblasts after NaCl or abscisic acid (ABA) treatment relative to the control (two replicates each). (C) log_2_ fold change after *RHD6* induction under dual stress (this study) and in public comparisons of low Pi or NaCl with the respective control in wild-type roots (datasets in Dataset S1). Grey cells with a line, not expressed (below 2 CPM in both groups in C; below 5 CPM in B); crosses, genes absent from the deposited table. (D) Spearman correlation between the log_2_ fold changes after *RHD6* induction and those of each public comparison (same columns as in C), across the differentially expressed genes of this study. (E) Co-expression with *RHD6* in low-Pi and NaCl datasets: Pearson correlations of expression standardized within each study, combined across studies, with 95% confidence intervals. n.e., not estimable (below 1 CPM in all or all but one of the studies). All values are given in Dataset S1. Alt text: Tracks of chromatin accessibility and H3K27me3 along the HEC1, HEC3 and EIL4 genes with the positions of the predicted binding motifs, a dot plot of expression in root cell types, heatmaps of the responses of selected genes to low phosphate or salt, a bar chart of correlations with the response to RHD6 induction, and a plot of co-expression with RHD6 in low-phosphate and salt datasets.

### Concluding remarks

Induced expression of *RHD6* keeps RHs growing under a combination of Pi deficiency and salinity that inhibits RH growth in the wild type. The transcriptome and the 396-gene core GRN provide a resource of candidate RHD6 targets under dual stress, and the mutant analysis identifies HEC3 as a modulator of the RH response. The study has clear limits: single-stress data only from public datasets, a single time point after 3 d of induction, whole-root samples, no β-estradiol control for the transcriptome and predicted rather than measured binding sites. Time-resolved and cell type-specific profiling, together with genome-wide binding data for RHD6, will be needed to separate direct from indirect targets. Because the function of class I *RSL* genes is conserved in grasses (Kim and Dolan, 2016), adjusting the dosage of RHD6 or of its downstream modulators could be explored as a way to maintain RH-mediated nutrient capture in saline, Pi-poor soils.

## Supplementary data

The following supplementary data are available at *JXB* online.

**Fig. S1**. Design of the TRANSPLANTA screen, identity of the transcription factor lines shown in Fig. 1A, and root hair length at increasing concentrations of β-estradiol.

**Fig. S2**. The core RHD6 gene regulatory network of 396 candidate direct targets grouped by functional class.

**Fig. S3**. T-DNA insertion sites and genotyping primers of the *hec1*, *hec3* and *eil4* mutants.

**Dataset S1**. TRANSPLANTA lines and root hair data of the screen; T-DNA insertion lines and genotyping primers; RNA-seq, promoter motif, co-expression network, statistical and public-data datasets; core network genes with the largest changes in expression after *RHD6* induction.

## Acknowledgements

We thank the TRANSPLANTA Consortium and the stock centre that supplied the TPT and T-DNA insertion lines. We thank Marek Mutwil and colleagues for providing the TEA-GCN co-expression data. Generative AI was used in the preparation of this work: Claude (Anthropic; Claude Fable 5.2) assisted with the statistical analysis and plotting of root hair data (Figs 1A, E, H and 2E, F), the Gene Ontology and promoter motif enrichment tests, the functional classification and null model of the core network, the re-analysis of public datasets (Fig. 3), the preparation of Figs 1, 2, 3, S2 and S3 and of Dataset S1, literature searches, the checking of references and the editing of the text. The authors directed this work, reviewed and verified all outputs and take full responsibility for the content.

## Author contributions

JP-D and MAI: investigation, formal analysis, writing – review & editing; PT-R and CM: formal analysis, data curation, writing – review & editing; FG, HS-G, GP and RA: formal analysis, writing – review & editing; JME: conceptualization, formal analysis, supervision, funding acquisition, writing – original draft. All authors read and approved the manuscript.

## Data availability

The RNA-seq data have been deposited at the NCBI under BioProject accession number PRJNA1565760. All other data supporting the findings of this study are available within the paper and its supplementary data. The public datasets re-analysed here are listed with their accession numbers in Dataset S1.

## Abbreviations

bHLH: basic helix–loop–helix;
CPM: counts per million;
DEG: differentially expressed gene;
FDR: false discovery rate;
GO: Gene Ontology;
GRN: gene regulatory network;
MS: Murashige and Skoog medium;
Pi: inorganic phosphate;
RH: root hair;
RNA-seq: RNA sequencing;
TF: transcription factor;
TPT: TRANSPLANTA.

